# The influence of biological and statistical properties of CpGs on epigenetic predictions of eighteen traits

**DOI:** 10.1101/2022.02.08.479569

**Authors:** Robert F. Hillary, Daniel L. McCartney, Allan F. McRae, Archie Campbell, Rosie M. Walker, Caroline Hayward, Steve Horvath, David J. Porteous, Kathryn L. Evans, Riccardo E. Marioni

## Abstract

**Background:** CpG methylation levels can help to explain inter-individual differences in phenotypic traits. Few studies have explored whether identifying CpG subsets based on biological and statistical properties can maximise predictions while minimising array content.

**Methods:** Variance component analyses and penalised regression (epigenetic predictors) were used to test the influence of (i) the number of CpGs considered, (ii) mean CpG methylation variability and (iii) methylation QTL status on the variance captured in eighteen traits by blood DNA methylation. Training and test sets comprised ≤4,450 and ≤2,578 unrelated individuals from Generation Scotland, respectively.

**Results:** As the number of CpG sites under consideration decreased, so too did the estimates from the variance components and prediction analyses. Methylation QTL status and mean CpG variability did not influence variance components. However, relative effect sizes were 15% larger for epigenetic predictors based on CpGs with methylation QTLs compared to sites without methylation QTLs. Relative effect sizes were 45% larger for predictors based on CpGs with mean beta-values between 10%-90% compared to those using hypo- or hypermethylated CpGs (beta-value ≤10% or ≥90%).

**Conclusion:** Arrays with fewer CpGs could reduce costs, leading to increased sample sizes for analyses. Our results show that reducing array content can restrict prediction metrics and careful attention must be given to the biological and distribution properties of CpGs in array content selection.

## Background

DNA methylation (DNAm) involves the addition of methyl groups to the fifth carbon of cytosine bases, typically in the context of cytosine-guanine dinucleotides (CpG sites). There are approximately 28 million CpG sites across the human genome (1, 2), of which 60-80% are methylated (3). Illumina DNAm arrays are popular technologies for profiling genome-wide DNAm at expert-selected subsets of CpG sites. The Infinium HumanMethylation 450K and HumanMethylationEPIC (EPIC) arrays cover 99% of RefSeq genes and interrogate 485,577 and 863,904 CpG sites, respectively (4, 5). DNAm data are routinely utilised in health outcomes research. First, the arrays are employed in association studies to uncover individual genomic loci associated with disease states and other phenotypes (6). Second, the total array content can be used to estimate the contribution of DNAm to inter-individual variability in human traits (7, 8). Third, machine learning algorithms can be applied to DNAm data to identify weighted linear combinations of CpG sites that predict numerous phenotypes, including chronological age, smoking status and body mass index (9–12).

Genetic, demographic, and environmental factors contribute to inter-individual variability in CpG methylation (13). Common genetic factors that correlate with CpG methylation are termed methylation quantitative trait loci (mQTLs) and explain on average 15% of the additive genetic variance of DNAm (14). Variation in CpG methylation might also reflect technical artefacts, including heterogeneity in sample preparation and batch effects (15). A large number of CpG sites exhibit low levels of inter-individual variation in a given tissue, such as blood (16–19). Several methods have been proposed to remove sites that are non-variable in a given tissue. The methods include mixture modelling, principal component analyses and empirically-derived data reduction strategies (20–22). In the context of locus discovery, these methods reduce the severity of multiple testing correction and might improve power to detect epigenetic associations with phenotypes. However, it is unclear if low-variability CpG sites affect the amount of phenotypic variance captured by DNAm. Lacking also are studies that examine the influence of CpG distribution properties on DNAm-based predictors.

CpG sites with high inter-individual variation in DNA methylation might be more informative for capturing variance in human traits compared to those that are invariant (i.e. low inter-individual variation). Here we tested the hypothesis that subsets of CpG sites that exclude invariant probes show similar predictive capacities for human traits as all available CpGs. We utilised blood DNAm data and eighteen phenotypes from 4,450 unrelated volunteers in the population-based cohort Generation Scotland (23, 24). We compared the performance of five primary sets of CpGs. The first set of CpGs included all sites common to the 450k and EPIC arrays (the reference set). In the second set, we excluded invariant CpGs (e.g. mean methylation signal (β) ≤10% or ≥90% across individuals) as well as those that are under genetic control (mQTLs) (14). The exclusion criteria allowed us to retain variable CpG sites and those whose variability might reflect environmental contributions. The third, fourth and fifth sets included the 10,000, 20,000 and 50,000 most variable non-mQTL CpGs (i.e. highest standard deviations (SDs)).

We used OmicS-data-based Complex trait Analysis (OSCA) to estimate the proportion of variability in eighteen lifestyle, physical and biochemical traits captured by a given CpG set (7). We also used penalised regression models to build DNAm-based predictors of the eighteen traits. We compared results from the five primary CpGs sets, which had decreasing numbers of CpGs and increasing mean CpG variabilities. As further analyses, we also considered subsets of CpGs with (i) an mQTL (with a mean β between 10% and 90%), (ii) hypo- or hypermethylated CpGs (with mean β≤10% or ≥90%) and (iii) genome-wide significant EWAS Catalog CpGs (at P<3.6 x 10^-8^). We compared results from these CpG sets against one another and the primary CpG sets, as well as against randomly sampled CpGs with equal CpG number.

## Methods

### Study Cohort

Details of Generation Scotland (GS) have been described previously (23, 24). GS is a family-based, genetic epidemiology cohort that consists of 24,084 volunteers. There were 5,573 families with a median size of 3 members (interquartile range = 2-5 members, excluding 1,400 singletons). Genome-wide DNAm was profiled using blood samples from GS baseline (2006-2011). DNAm was processed in two separate sets of 5,200 (2016) and 4,585 samples (2019) (25). The sets are hereafter referred to as ‘Set 1’ and ‘Set 2’, respectively.

### Preparation of DNA methylation data

DNAm was measured using the Infinium MethylationEPIC BeadChip at the Wellcome Clinical Research Facility, Western General Hospital, Edinburgh. Methylation typing in Set 1 (n=5,200) and Set 2 (n=4,585) was performed using 31 batches each. Full details on the processing of DNAm data are available in Additional File 1. Poor-performing and sex chromosome probes were excluded, leaving 760,943 and 758,332 CpGs in Set 1 and 2, respectively. Participants with unreliable self-report questionnaire data (self-reported positive for 20 diseases in the questionnaire), saliva samples and possible XXY genotypes were excluded, leaving 5,087 and 4,450 samples in Set 1 and 2, respectively. In Set 1, there were 2,578 unrelated individuals (common SNP GRM-based relatedness <0.05). In Set 2, all 4,450 individuals were unrelated to one another. Individuals in Set 1 were unrelated to those in Set 2. Set 2 was used for OSCA models and as the training sample in DNAm-based prediction analyses given its larger sample size (n=4,450). Set 1 was used as the test set in DNAm-based prediction analyses (n=2,578). Linear regression models were used to correct CpG β values for age, sex and batch effects separately in Set 1 (*test set*, n=2,578) and Set 2 (*training set*, n=4,450).

### Identification of variable blood CpG sites

There were 758,332 CpG sites in Set 2 following quality control. First, we restricted CpG sites to those that are common to the 450k and EPIC arrays to allow for generalisability to other epigenetic studies (n=398,624 CpGs). We excluded CpGs that were predicted to cross-hybridise and those with polymorphisms at the target site, which can alter probe binding (n_CpGs_=4,970 excluded, 393,654 remaining) (26, 27). These 393,654 CpGs represented the set termed ‘all available CpGs’ in our analyses.

We defined a set of criteria to identify variable blood CpG sites. First, we removed CpG sites that are hypo- or hypermethylated in the sample (i.e. mean methylation β value ≤10% or ≥90%, respectively, n_CpGs_=144,150 excluded). Hypo- and hypermethylated CpGs had a mean SD of 0.01 (range=0.002-0.13). CpGs with mean β between 10% and 90% (n_CpGs_=249,504) had a mean SD of 0.03 (range=0.008-0.33). Second, we excluded 133,758 CpGs that overlapped with known blood-based mQTLs (GoDMC (28), P value<5×10^-8^). There were 115,746 remaining sites, which represented the ‘variable non-mQTL CpGs’ subset. We then extracted the 10,000, 20,000 and 50,000 non-mQTL CpGs with the highest SDs (Additional File 2: Table S1).

### Preparation of phenotypic data

Eighteen traits were considered in our analyses. These were chronological age, seven biochemical traits (creatinine, glucose, high-density lipoprotein cholesterol, potassium, sodium, total cholesterol and urea) and ten complex traits (alcohol consumption, body fat percentage, body mass index, diastolic blood pressure, forced expiratory volume in one second (FEV), forced vital capacity (FVC), heart rate (average beats/minute), smoking pack years, systolic blood pressure, waist-to-hip ratio). Full details on phenotype preparation are detailed in Additional File 1.

The seventeen biochemical and complex traits were trimmed for outliers (i.e. values that were ± 4 SDs away from the mean). Fifteen phenotypes (excluding FEV and FVC) were regressed on age, age-squared and sex. FEV and FVC were regressed on age, age-squared, sex and height (in cm). Correlation structures for raw (i.e. unadjusted) and residualised phenotypes are shown in Additional File 3: Figure S1-S2, respectively. For age models, DNAm and chronological age (in years) were unadjusted. Residualised phenotypes were entered as dependent variables in OSCA or penalised regression models.

### Variance component analyses

OSCA software was used to estimate the proportion of phenotypic variance in eighteen traits captured by DNAm (7). Omic-data-based relationship matrices (ORMs) were generated for all CpG sets. Restricted maximum likelihood (REML) estimated the proportion of phenotypic variance captured by CpGs that were used to build a given ORM.

### LASSO regression and prediction analyses

Least absolute shrinkage and selector operator (LASSO) regression was used to build DNAm-based predictors of eighteen phenotypes. The R package *biglasso* (29) was implemented and the training sample included ≤4,450 samples from Set 2. The mixing parameter (alpha) was set to 1 and tenfold cross-validation was applied. The model with the lambda value that corresponded to the minimum mean cross-validated error was selected. Epigenetic scores for traits were derived by applying coefficients from this model to corresponding CpG sites in the test set (n=2,578).

Linear regression models were used to test for associations between DNAm-based predictors (i.e. epigenetic scores) for the eighteen traits and their corresponding phenotypic values in Set 1. The incremental r-squared (R^2^) was calculated by subtracting the R^2^ of the full model from that of the null model (shown below). For the FEV and FVC predictors, height was included as an additional covariate in both models. For the age predictors, the R^2^ value pertained to that of the epigenetic score without further covariates.

*Null model*: Phenotype ~ chronological age + sex
*Full model*: Phenotype ~ chronological age + sex + epigenetic score

### Sub-sampling analyses

We tested whether variance components and incremental R^2^ estimates from CpG sets were significantly different from those expected by chance. For OSCA estimates, we generated 1,000 sub-samples of 10,000, 20,000, 50,000 and 115,746 CpGs (to match the ‘variable non-mQTL CpGs’ set). We also generated 100 sub-samples of 10,000, 20,000, 50,000 and 115,746 CpG sets for LASSO regression to lessen the computational burden. The sub-sampled CpG sets were derived from ‘all available CpGs’ (n_CpGs_=393,654).

We tested whether the most variable CpGs were significantly over-represented or under-represented for genomic and epigenomic annotations. The annotations were derived from the *IlluminaHumanMethylationEPICanno.ilm10b4.hg19* package in R (30). Annotations for the 5,000 and 10,000 most variable non-mQTL CpGs were compared against 1,000 sub-samples of non-mQTL CpGs with equal CpG number. Here CpGs were sub-sampled from the ‘variable non-mQTL CpGs’ set (n_CpGs_=115,746) and not from ‘all available CpGs’ (n_CpGs_=393,654) as the latter contains CpGs with and without mQTLs, which show different genetic architectures (28).

### Comparisons of methylation QTL status and mean methylation beta-value levels

In addition to non-mQTL CpG subsets (with mean β between 10% and 90%), we tested two further classes of CpG. First, we considered CpGs with an mQTL from GoDMC (P<5×10^-8^) that had mean β between 10% and 90% (n_CpGs_=133,758) (14). Second, we considered all hypo- or hypermethylated CpGs (β≤10% or ≥90%, n_CpGs_=144,150). We tested the performances of the 10,000, 20,000, 50,000, and 115,746 most variable CpGs from each of these three CpG classes.

We also repeated REML and LASSO regression using EWAS Catalog CpGs (31). EWAS Catalog CpGs contained CpGs with an mQTL, CpGs without an mQTL and hypo- and hypermethylated CpGs. We restricted EWAS Catalog CpGs to those with P<3.6 x 10^-8^ (32) and those reported in studies with sample sizes >1,000. We also excluded studies related to chronological age due to the very large number of sites implicated, and those in which Generation Scotland contributed to analyses. There were 100 studies that passed inclusion criteria with 47,093 unique CpGs. Of these, 38,853 CpGs overlapped with ‘all available CpGs’ used in our analyses (n_CpGs_=393,654). To allow for comparison to other CpG subsets, the 10,000 and 20,000 most variable EWAS Catalog CpGs (n_CpGs_=38,853) were extracted.

## Results

Demographics and summary data for all phenotypes are shown in Additional File 2: Table S2. The mean age in Set 1 was 50.0 years (SD = 12.5) and the sample was 61.4% female. Set 2 showed a similar mean age of 51.4 years (SD = 13.2) with a slightly lower proportion of females (56.3%). Values for all other phenotypes were comparable between the sets.

### Phenotypic variance captured by DNAm decreases with the number of CpGs considered

We compared variance component estimates from ‘all available CpGs’ (n_CpGs_=393,654) with four subsets of CpGs containing 10,000, 20,000, 50,000 and 115,746 sites. The subsets contained CpGs with mean β between 10% and 90% and without underlying mQTLs (i.e. non-mQTL CpGs) (Figure 1). The subset that contained 115,746 CpGs represented all non-mQTL CpGs with mean β between 10% and 90% (i.e. ‘variable non-mQTL CpGs’). The remaining three CpG subsets harboured the 10,000, 20,000 and 50,000 most variable of these sites, showing the highest standard deviations in Set 2 (n=4,450).

**Figure 1.**
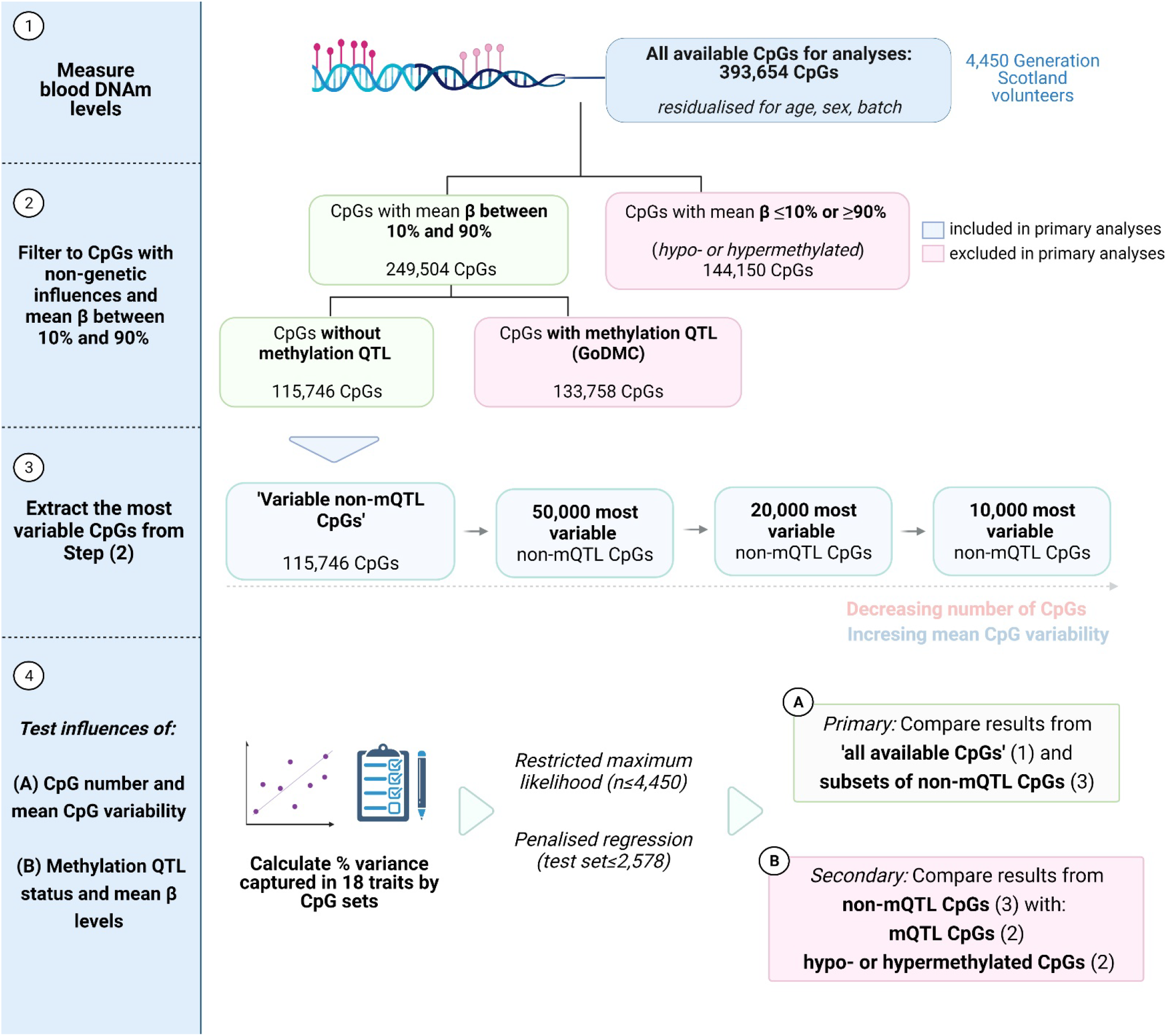
Overview of analysis strategy to test the influences of the number of CpGs considered, CpG distribution properties and methylation QTL status on epigenetic predictions of eighteen traits. We tested whether subsets of CpGs showed similar predictive capacities to total DNAm array content (1) (‘all available CpGs’, n=393,654). We first identified CpG subsets of interest. We restricted primary analyses to CpGs without genetic influences (i.e. non-mQTL CpGs) and those with mean beta-values (β) between 10% and 90% (2). These CpGs were termed ‘variable non-mQTL CpGs’ (n=115,746). We then extracted the 10,000, 20,000 and 50,000 CpGs with the highest standard deviations from the pool of 115,746 non-mQTL CpGs (3). In our primary analyses, we compared the predictive performances of these four CpG subsets against that of the full set of CpGs used in our analyses (4). In further analyses, we tested the relative performances of CpG subsets based on (i) CpGs without mQTLs and with mean beta between 10%-90% (shown in green in (2), highlighted in (3)), (ii) CpGs with mQTLs and with mean beta between 10%-90% (shown in red in (2)) and hypo- or hypermethylated CpGs (mean beta ≤10% or ≥90%, also shown in red in (2)). DNAm, DNA methylation; mQTL, methylation quantitative trait locus; SD, standard deviation. Image created using Biorender.com.

The proportion of phenotypic variance captured by ‘all available CpGs’ (n_CpGs_ = 393,654) ranged from 23.7% (standard error (se) = 6.0%) for blood potassium levels to 79.6% (se=2.1%) for smoking pack years (Additional File 2: Table S3). The average proportion of variance captured across seventeen biochemical and complex traits was 54.0%. Mean estimates were 44.1% and 61.0% for biochemical and complex traits, respectively (Additional File 3: Figure S3).

The four CpG subsets (in order of increasing size) on average captured 21.9%, 30.4%, 40.6% and 47.9% of phenotypic variance across seventeen traits (Additional File 2: Table S3). Generally, the estimates were not significantly different from sub-sampled CpG subsets of equal size (Additional File 2: Table S4). An exception to this was smoking pack years (P<0.05). Figure 2 shows the four traits with the highest proportions of phenotypic variance captured by CpG methylation.

**Figure 2.**
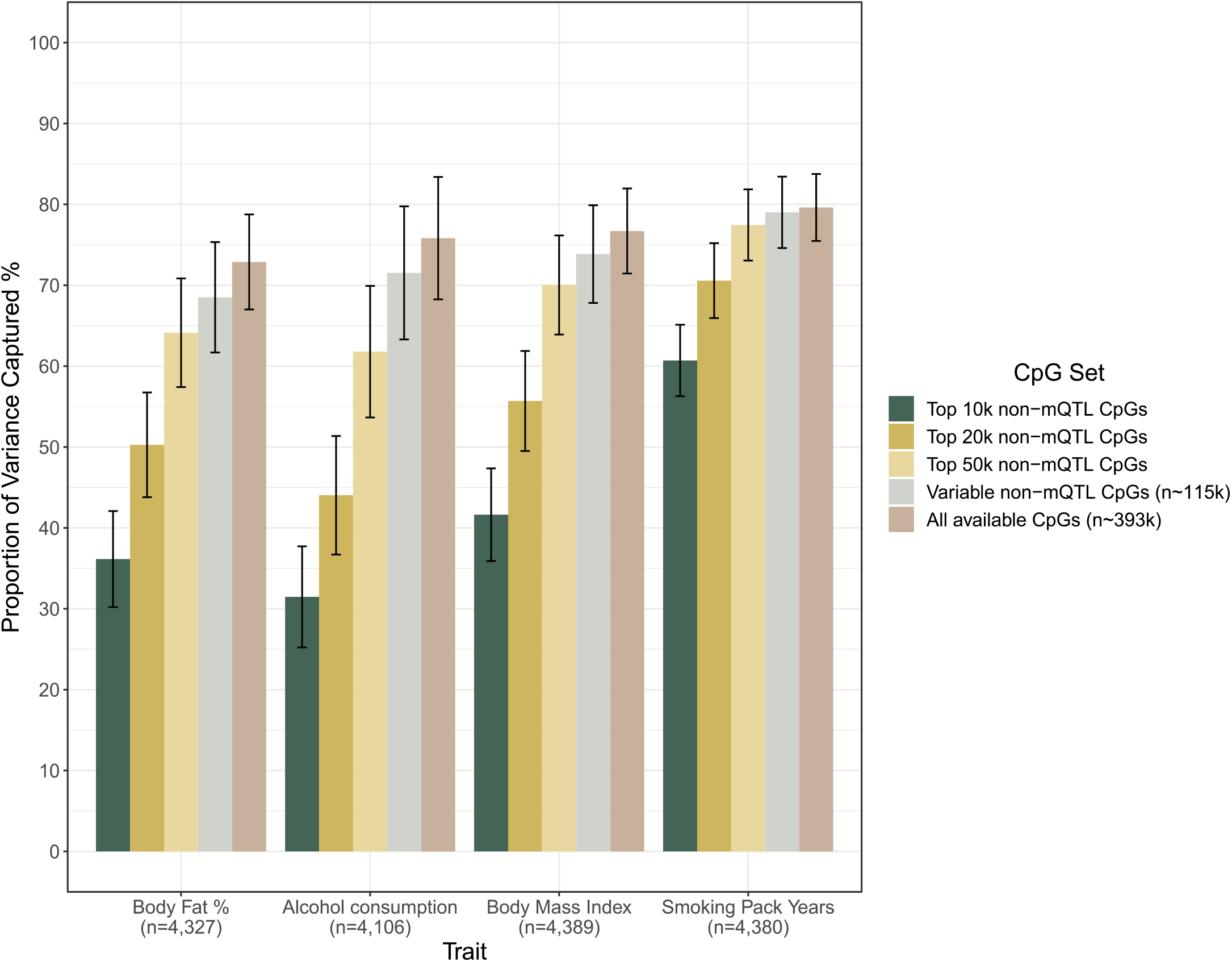
Phenotypic variance captured in complex traits by four CpG subsets of increasing size and all available CpGs. Restricted maximum likelihood was used to estimate variance components in Set 2 of Generation Scotland (n ≤4,450, OSCA software). The four traits (out of seventeen biochemical and complex traits) with the highest proportion of variance captured by DNAm are shown. The four traits were: body fat percentage (%, n=4,327), alcohol consumption (units/week, n=4,106), body mass index (kg/m^2^, n=4,389) and smoking pack years (n=4,380). Five different sets of CpGs were compared. ‘All available CpGs’ denotes CpGs that were common to the Illumina EPIC and 450k arrays and passed quality control procedures in Set 2 of Generation Scotland (n=393,654 CpGs). The ‘variable non-mQTL CpGs’ set consisted of CpGs with non-genetic influences and mean beta-values between 10% and 90%. The remaining three CpG subsets contained the 10,000, 20,000 and 50,000 most variable non-mQTL CpGs (ranked by their standard deviations). Vertical bars show 95% confidence intervals. DNAm, DNA methylation; mQTL, methylation quantitative trait locus; OSCA, OmicS data-based Complex Trait Analysis.

### Performance of DNAm-based predictors decreases with the number of CpGs considered

DNAm-based predictors based on ‘all available CpGs’ (n_CpGs_=393,654) captured between 0.74% (FVC) and 46.0% (smoking pack years) of trait variance in the test set (Additional File 2: Table S5). DNAm-based predictors developed from ‘all available CpGs’ on average captured 9.1% of trait variance (Additional File 3: Figure S4).

DNAm-based predictors developed from the four subsets of non-mQTL CpGs (in order of increasing size) captured 5.0%, 5.6%, 6.6% and 6.7% of phenotypic variation. The four traits with the highest incremental R^2^ estimates are shown in Figure 3.

**Figure 3.**
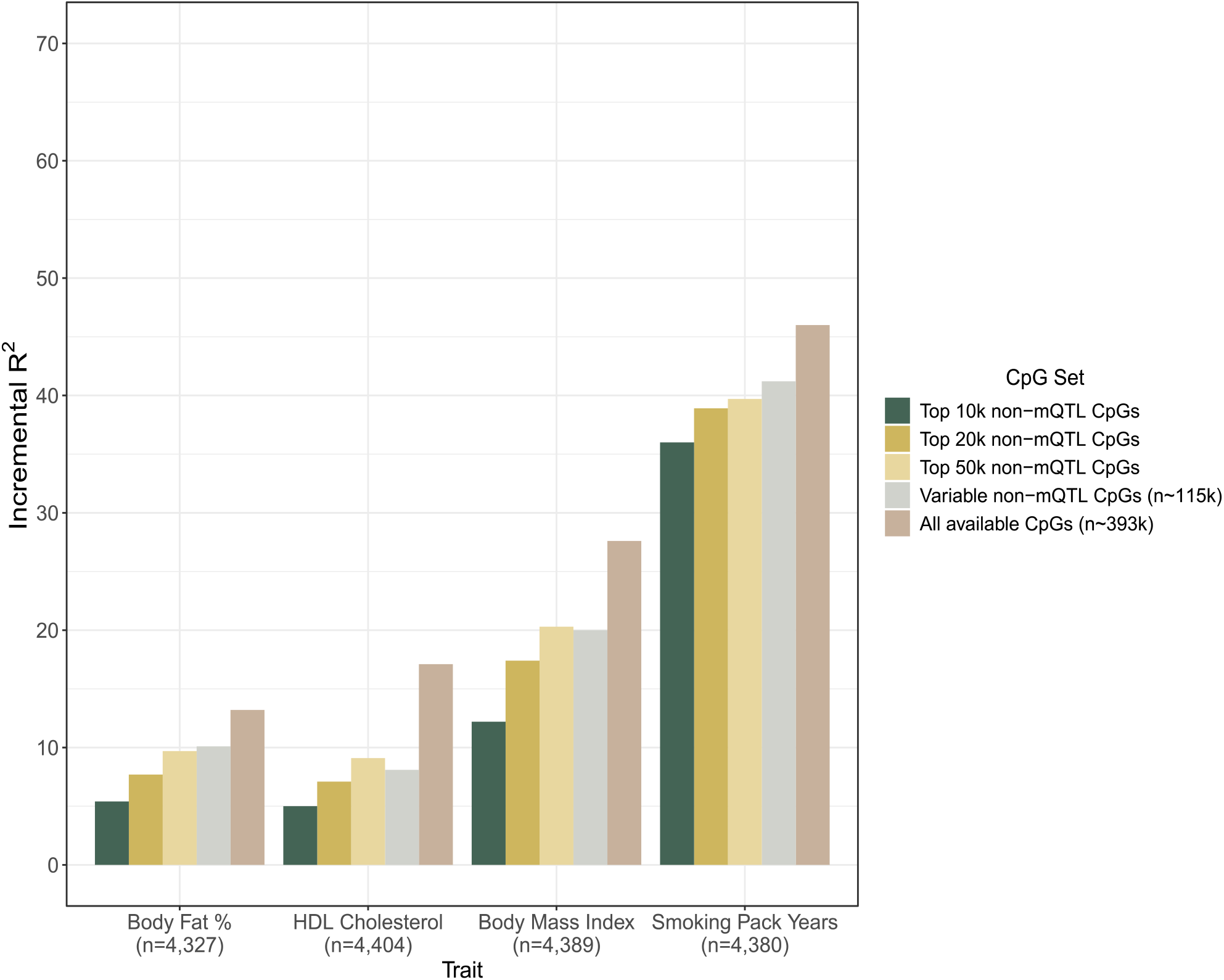
DNAm-based prediction of complex traits using four CpG subsets of increasing size and all available CpGs. LASSO regression was used to build blood DNAm-based predictors of seventeen biochemical and complex traits (n=4,450 Generation Scotland samples in training set). An unrelated sample of 2,578 individuals in Generation Scotland served as the test set. The four traits with the highest proportion of variance captured by DNAm predictors are displayed (incremental R^2^ estimates above null model, see main text). The four traits were: body fat percentage (%, n=4,327), HDL cholesterol (mmol/L, n=4,404), body mass index (kg/m^2^, n=4,389) and smoking pack years (n=4,380). The first set of CpGs included those that passed quality control in the training sample, were common to both the EPIC and 450k arrays and included both CpGs with methylation QTLs (mQTLs) and CpGs without mQTLs. The next four sets of CpGs included non-mQTL CpGs only and had decreasing numbers of CpGs but increasing mean CpG variabilities. DNAm, DNA methylation; HDL, high-density lipoprotein; LASSO, Least Absolute Shrinkage and Selection Operator; mQTL, methylation quantitative trait locus.

The performances of the four subsets of non-mQTL CpGs were weaker for biochemical measures than complex traits. For biochemical measures, their effect sizes were 19.1%-38.7% of the magnitude of estimates from ‘all available CpGs’. The corresponding estimates were 47.5%-74.2% for complex traits (Additional File 2: Table S5). Incremental R^2^ estimates from the four CpG subsets were not significantly different from sub-sampled CpG sets of equal size (Additional File 2: Table S6).

### Subsets of CpGs capture similar amounts of variation in chronological age as all available CpGs

Using REML, ‘all available CpGs’ captured 100% of variability in chronological age (n_CpGs_=393,654). CpG subsets containing 20,000, 50,000 and 115,746 CpGs with non-genetic influences also captured 100% of the variance. The subset containing the 10,000 most variable non-mQTL CpGs captured 92.1% (se=0.9%, Additional File 2: Table S7).

An epigenetic age predictor based on ‘all available CpGs’ captured 91.7% of the variance in chronological age (n=2,578). The R^2^ estimates from four CpG subsets containing 10,000, 20,000, 50,000 and 115,746 CpGs were 83.9%, 85.7%, 87.7%, 87.4%, respectively (Additional File 2: Table S8). The estimates were not significantly different from those in randomly sampled subsets with an equivalent number of CpG sites.

### Highly variable CpGs are enriched for intergenic and upstream features

The 5,000 most variable CpGs (without an mQTL) were over-represented in intergenic sites (fold enrichment [FE]=1.2, P<0.001) and sites 200-1,500 bases upstream from the transcription start site (TSS1500, FE=1.1, P=0.02). The 5,000 most variable CpGs were significantly under-represented in 3’UTR regions (FE=0.87, P=0.004), gene body CpGs (FE=0.86, P=0.001) and CpGs 2-4 kilobases downstream (3’) from a CpG island (Southern Shelf, FE=0.89, P = 0.01). These patterns were also present for the 10,000 most variable non-mQTL CpGs (Additional File 2: Table S9).

### Methylation QTL status and mean methylation beta-value levels do not influence variance component estimates

We performed further analyses to determine the relative predictive capacities of four subsets, or classes, of CpGs. The first three subsets were: (i) CpGs *without* an mQTL and mean β between 10% and 90% (considered in primary analyses), (ii) CpGs *with* an mQTL and mean β between 10% and 90% and (iii) CpGs with mean β ≤10% or ≥90%, that is, hypo- or hypermethylated CpGs. The latter two classes are shown as the excluded CpGs in Figure 1. We also considered a fourth class, which was EWAS Catalog CpGs (n=38,853, see Methods). The EWAS Catalog CpGs contained all three of the other classes: >65% were CpGs with an mQTL and <5% were hypo- or hypermethylated (Additional File 2: Table S1).

Across all classes, variance estimates decreased with the number of CpGs under consideration (Table 1). All CpG classes, when matched for CpG number, showed comparable variance component estimates (Table 1, Additional File 2: Table S10-S12). An exception to this involved subsets that included 115,746 CpGs. CpGs with mean β between 10% and 90% on average captured 10% more trait variance than hypo- or hypermethylated CpGs (β≤10% or ≥90%) at this threshold. The CpG classes captured similar amounts of variance in age (Additional File 2: Table 13).

**Table 1.** The influences of the number of CpGs, mean CpG variability and methylation QTL status on variance component estimates.

| CpG Classification | n<sub>CpG</sub> | 10,000 | 20,000 | 50,000 | 115,746 |
| --- | --- | --- | --- | --- | --- |
| CpGs without an mQTL | 115,746 | 21.9% | 30.4% | 40.6% | 47.9% |
| CpGs with an mQTL | 144,150 | 16.5% | 26.1% | 38.7% | 48.0% |
| Hypo- or hypermethylated CpGs | 133,758 | 18.4% | 27.7% | 37.9% | 38.7% |
| EWAS Catalog CpGs | 38,853 | 24.4% | 32.9% | - | - |

Metric shown is the average % of variance captured in seventeen biochemical and complex traits. mQTL, methylation QTL.

### CpGs with methylation QTLs and intermediate methylation levels are important for out-of-sample trait predictions

Epigenetic predictors based on EWAS Catalog CpGs (n=38,853) captured as much variance as those based on ‘all available CpGs’ (n_CpGs_=393,654). The 10,000 and 20,000 most variable EWAS Catalog CpGs showed estimates that were 85.3% and 91.5% of the magnitude of those from all ‘available CpGs’ (Additional File 2: Table S14-S16).

Epigenetic predictors based on CpGs with an mQTL (n=133,758), and the 115,746 most variable of these CpGs, also captured as much phenotypic variance as predictors based on ‘all available CpGs’ (Additional File 2: Table S14). Exceptions included predictors for creatinine and systolic blood pressure (60-70% of estimates from ‘all available CpGs’).

The relative effect sizes (i.e. relative incremental R^2^ estimates) were on average 15% larger for CpGs with mQTLs versus non-mQTL CpGs. Relative effect sizes were also approximately 45% greater for CpGs with mean β between 10% and 90% when compared to hypo- or hypermethylated CpGs with mean β ≤10% or ≥90% (Table 2, Additional File 2: Table S14-S16).

**Table 2.** The influences of the number of CpGs, mean CpG variability and methylation QTL status on DNAm-based predictions.

| CpG Classification | n<sub>CpG</sub> | 10,000 | 20,000 | 50,000 | 115,746 |
| --- | --- | --- | --- | --- | --- |
| CpGs without an mQTL | 115,746 | 5.0% | 5.6% | 6.6% | 6.7% |
| CpGs with an mQTL | 144,150 | 4.8% | 6.2% | 8.1% | 9.0% |
| Hypo- and<br>hypermethylated CpGs | 133,758 | 2.8% | 3.3% | 4.0% | 4.2% |
| EWAS Catalog CpGs | 38,853 | 8.0% | 8.7% | - | - |

Metric shown is the average % of variance captured in seventeen biochemical and complex traits by DNAm-based predictors. mQTL, methylation QTL.

The performances of age predictors were comparable for all CpG classes except hypo- and hypermethylated CpGs, which showed R^2^ estimates that were 5%-10% lower than other CpG classes (Additional File 2: Table S17).

## Discussion

The amount of phenotypic variance captured by DNAm decreased in all traits as the number of CpGs under consideration decreased. Further, variance component estimates were similar for CpG subsets with and without genetic influences and CpG subsets with and without hypo- and hypermethylated CpGs. The estimates were also comparable to sub-sampled CpG subsets of equal size. Therefore, the number of CpGs considered is an important determinant of the amount of within-sample trait variance that can be captured by DNAm. Methylation QTL status and mean beta-value levels did not appear to impact variance component estimates. By contrast, epigenetic predictors based on CpG subsets with mQTLs generally outperformed those that contained CpGs without underlying mQTLs. Similarly, CpGs that had mean β between 10% and 90% outperformed subsets that contained hypo- and hypermethylated CpGs in out-of-sample trait predictions. Therefore, methylation QTL status and mean methylation levels are important factors in the performance of epigenetic trait predictions. As with variance component analyses, decreasing the number of CpGs considered resulted in poorer performing epigenetic predictors.

Highly variable CpGs were enriched for intergenic sites, which is consistent with the existing literature (19, 33, 34). However, the most variable CpGs that fall outside of CpG islands can be poorly captured by arrays (35). The list of the most variable CpGs might show variation between epigenomic datasets given differences in normalisation methods and systematic differences in cohort profiles. We also did not correct for additional covariates, such as cell-type heterogeneity, which could lead to differences in estimates for CpG variabilities. However, OSCA can account for unmeasured confounders and correlation structures between distal probes induced by confounders (7). We selected standard deviations to measure variability in CpG methylation levels. However, some CpGs may show non-normal distributions of beta-values or multimodal distributions (such as mQTL CpGs). This complicates the general application of one measure of variability across all CpG sites. Nevertheless, our results showed comprehensively that decreasing the number of available CpG sites reduced variance estimates regardless of mQTL status or mean methylation intensity.

High R^2^ estimates from subsets based on EWAS Catalog CpGs likely reflects contributions from all CpG classes (i.e. CpGs with and without an mQTL and hypo- or hypermethylated CpGs) and that many of the traits considered in this study feature in the EWAS Catalog. The superior performance of epigenetic predictors from mQTL CpG subsets compared to non-mQTL CpG subsets likely reflects the exclusion of CpG sites with strong biological signals in the latter. For instance, cg06500161 (*ABCG1*) and cg05575921 (*AHRR*) have underlying mQTLs and are strong epigenetic correlates of body mass index and smoking pack years, respectively (11, 36–40). CpGs with an mQTL are more reliably measured than those without mQTLs (41). Further, the training and test sets show similar genetic backgrounds, which might have supported replication of associations between traits and predictors based on CpGs with genetic influences. The relative performances of the predictors should be tested in cohorts of different ethnicities and clinical populations.

Traits with strong epigenetic correlates were the most robust to changes in CpG class or the number of CpGs considered. For instance, 20,000 CpGs were sufficient to capture 100% of inter-individual variation in chronological age. Epigenetic predictors based on 10,000 CpGs (with mean β between 10% and 90%) were 90% as accurate as a predictor based on ‘all available CpGs’. Highly accurate epigenetic predictors of chronological and biological age continue to be described (9, 10, 42–45). We show that (i) small subsets of CpGs can capture age-related changes in DNAm, (ii) DNAm-based age predictors are not affected by mQTL status and (iii) CpGs that are hypo- or hypermethylated are less informative for predicting age than CpGs with β values between 10% and 90%.

## Conclusion

Restricting DNAm array probes to the most variable sites could improve power in association studies whilst minimising array content. We show that this approach hampers variance component analyses, and that phenotypes with strong epigenetic correlates are the most robust to changes in the number of available CpGs. Further, CpGs with an mQTL and CpGs with intermediate DNAm levels are central to epigenetic predictions of clinically-relevant phenotypes. Our results demonstrate that strategies aiming to minimise arrays using fewer CpGs must carefully select CpG content in order maximise epigenetic predictions of human traits.

## Ethics approval and consent to participate

All components of the Generation Scotland study received ethical approval from the NHS Tayside Committee on Medical Research Ethics (REC Reference Numbers: 05/S1401/89 and 10/S1402/20). All participants provided broad and enduring written informed consent for biomedical research. Generation Scotland has also been granted Research Tissue Bank status by the East of Scotland Research Ethics Service (REC Reference Number: 20-ES-0021). This study was performed in accordance with the Helsinki declaration.

## Acknowledgments

This research was funded in whole, or in part, by Wellcome [104036/Z/14/Z]. For the purpose of open access, the author has applied a CC BY public copyright licence to any Author Accepted Manuscript version arising from this submission. We are grateful to the families who took part in this study, the general practitioners and the Scottish School of Primary Care for their help in recruiting them and the wider Generation Scotland team. Generation Scotland received core support from the Chief Scientist Office of the Scottish Government Health Directorates [CZD/16/6] and the Scottish Funding Council [HR03006]. DNA methylation profiling of the Generation Scotland samples was carried out by the Genetics Core Laboratory at the University of Edinburgh Clinical Research Facility, Edinburgh, Scotland. The DNAm profiling and analysis was supported by a Wellcome Investigator Award [220857/Z/20/Z] and Grant [104036/Z/14/Z] (PI: AM McIntosh) and through funding from Brain & Behavior Research Foundation ([27404], awardee: Dr DM Howard) and the Royal College of Physicians of Edinburgh (Sim Fellowship; Awardee: Dr HC Whalley). D.J.P is supported by Wellcome as PI, and MRC and NIHR grants as co-PI, made to the University of Edinburgh. C.H. is supported by an MRC University Unit Programme Grant MC_UU_00007/10 (QTL in Health and Disease). K.L.E was supported by a grant from Alzheimer’s Research UK, paid to the University of Edinburgh. R.E.M. is supported by Alzheimer’s Society major project grant AS-PG-19b-010. R.F.H., S.H., and R.E.M. are supported by a National Institutes of Health U01 grant, U01AG060908–01.

## Data Availability

According to the terms of consent for Generation Scotland participants, access to data must be reviewed by the Generation Scotland Access Committee. Applications should be made to.

## Code Availability

All code is available at the following Github repository: https://github.com/robertfhillary/Biological_Statistical_Properties_CpGs_Epigenetic_Predictions.

