## Additional File 1 - Supplementary Methods for "The influence of biological and statistical properties of CpGs on epigenetic predictions of eighteen traits"

**Additional File 1 - Supplementary Methods.** The following information pertains to Supplementary Methods for the manuscript *'The influence of biological and statistical properties of CpGs on epigenetic predictions of eighteen traits'* by Hillary *et al.*

**Preparation of DNA methylation data**

In Set 1, the *ShinyMethyl* package in R was used to compare plots of log median signal intensities across methylated and unmethylated beads in each array (1). Outliers were removed based on a visual inspection of the plots. The *wateRmelon* package in R was used to remove samples based on the following exclusion criteria: (i) >1% of probes had a detection P value > 0.05, (ii) probes had a beadcount <3 in >5% of samples and (iii) probes were non-autosomal i.e. XY probes (2). Eighty samples and 5,910 probes were excluded based on these criteria. Probes that were predicted to have off-target effects were excluded. Probes were also removed if they contained a SNP in the final five 3' bases or at the site of single-base extension in type I probes (n = 84,352) (3, 4). Twelve individuals were removed due to discordance between their methylation-based predicted sex and recorded sex. Seven further samples were removed as they were identified as genetic outliers in principal component analyses of genotype data (5). Ten samples that were derived from saliva were excluded, along with three individuals that self-reported 'Yes' to all health conditions listed on study questionnaires at baseline. One individual was removed as their methylation data indicated that they might have an XXY genotype. Data were normalised using the dasen method in the *wateRmelon* package. The final set consisted of 5,087 participants and 760,943 loci (6). In Set 1 there are 2,578 unrelated individuals.

In Set 2, the *Meffil* package in R was used to perform initial quality control steps (7). Samples were excluded if they met the following criteria: (i) there was a mismatch between self-reported and methylation-based predicted sex, (ii) more than 1% of probes had a detection P value > 0.05, (iii) samples showed evidence of dye bias, (iv) sample were outliers at bisulfite conversion control probes and (v) the sample had a median methylated signal intensity that was ≥ 3 standard deviations lower than expected. *ShinyMethyl* was then used to exclude probes based on the same criteria as those applied in Set 1. MDS of methylation data was performed and outliers were removed based on visual inspection of the resultant plots. *Meffil* was employed again to remove poor-performing probes that met the following exclusion criteria: (i) probes had a beadcount of <3 in >5% of samples and (ii) >5% of samples had a detection P value > 0.05. In total, 8,878 poor-performing probes and 135 samples were excluded. Probes with potential off-target effects were excluded as in Set 1. Probes were also removed if they had a SNP in the final five 3’ bases or in the single-base extension site (n = 84,352) (3, 4). Data were normalised using the dasen method. There were 4,450 individuals in Set 2 and 758,332 CpG sites following quality control procedures. The individuals in Set 2 are unrelated to each other and to those in Set 1.

**Preparation of phenotypic data**

Eighteen traits were considered: (1) alcohol consumption (units/week), (2) chronological age (years), (3) creatinine (µmol/L), (4) body fat percentage (%), (5) body mass index (kg/m$^2$), (6) diastolic blood pressure (mmHg), (7) forced expiratory volume in one second (FEV; L), (8) forced vital capacity (L), (9) glucose (mmol/L), (10) heart rate (average beats/min), (11) high-density lipoprotein (HDL) cholesterol (mmol/L), (12) potassium (mmol/L), (13) smoking pack years, (14) sodium (mmol/L), (15) systolic blood pressure (mmHg), (16) total cholesterol (mmol/L), (17) urea (mmol/L) and (18) waist-to-hip ratio.

A number of phenotypes underwent additional pre-processing steps as those outlined in the main manuscript. To reduce skewness in the distribution of alcohol consumption, a log(units + 1) transformation was performed. Body mass index was trimmed for extreme values at <17 and >50 kg/m$^2$. Body mass index values were log transformed to reduce skewness. Phenotypes were pre-adjusted as detailed in the main manuscript. Smoking pack years were calculated by multiplying the number of packs of cigarettes smoked per day by the number of years the individual has smoked. To reduce skewness, a log(pack years + 1) transformation was applied to the smoking pack years variable.
