## Additional File 3 - Supplementary Figures for "The influence of biological and statistical properties of CpGs on epigenetic predictions of eighteen traits"

**Additional File 3 - Supplementary Figures.** The following information pertains to Supplementary Figures for the manuscript *'The influence of biological and statistical properties of CpGs on epigenetic predictions of eighteen traits'* by Hillary *et al.*

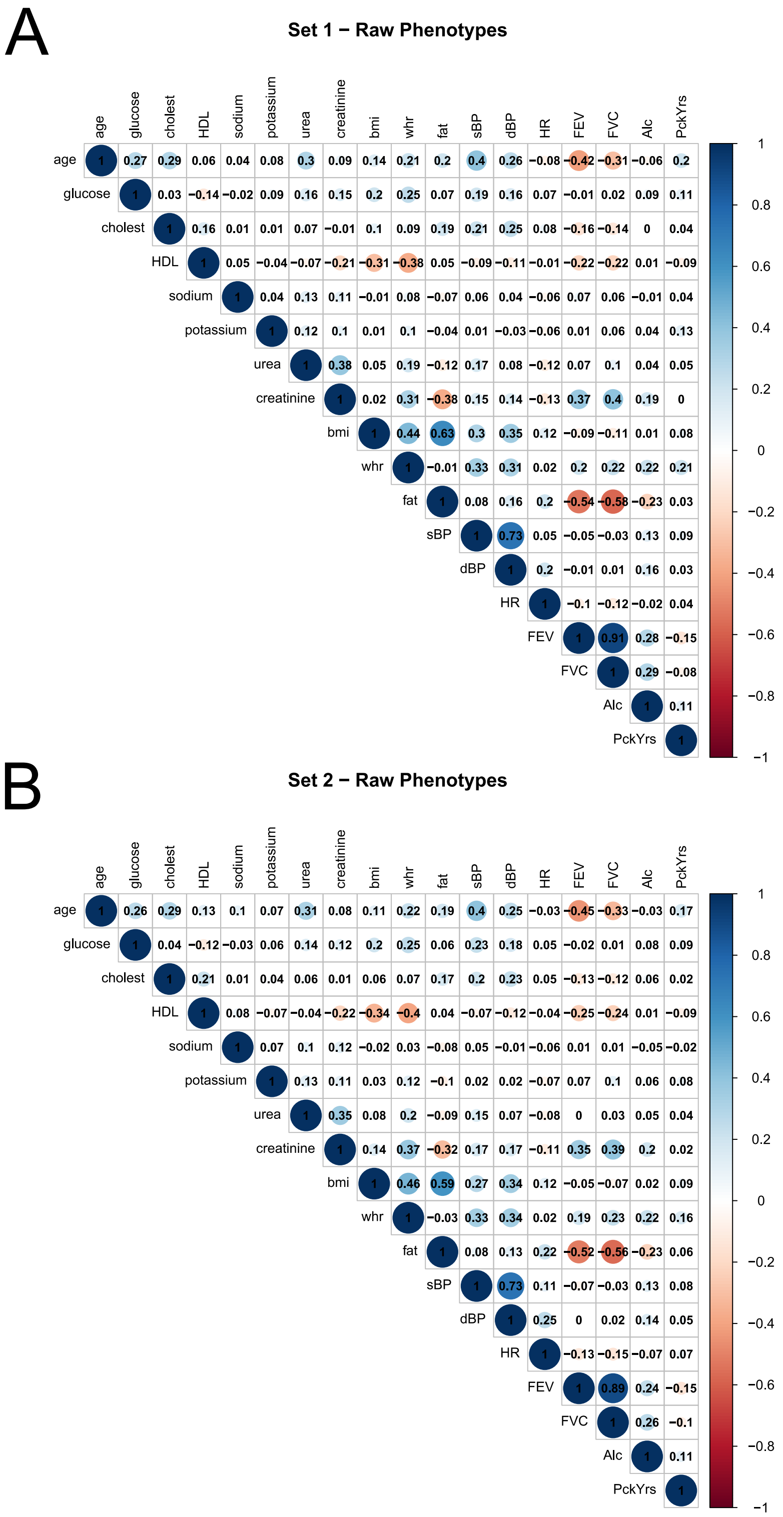


**Figure S1. Correlation structure between raw (i.e. unadjusted) phenotypes in Set 1 and Set 2 of Generation Scotland.** Set 1 (A) and Set 2 (B) had 2,578 and 4,450 unrelated individuals, respectively. Alc, alcohol consumption; bmi, body mass index; cholest, total cholesterol; dBP, diastolic blood pressure; fat, body fat percentage; FEV, forced expiratory volume in one second; FVC, forced vital capacity; HDL, high-density lipoprotein cholesterol; HR, heart rate; PckYrs, smoking pack years; sBP, systolic blood pressure; whr, waist-to-hip ratio.

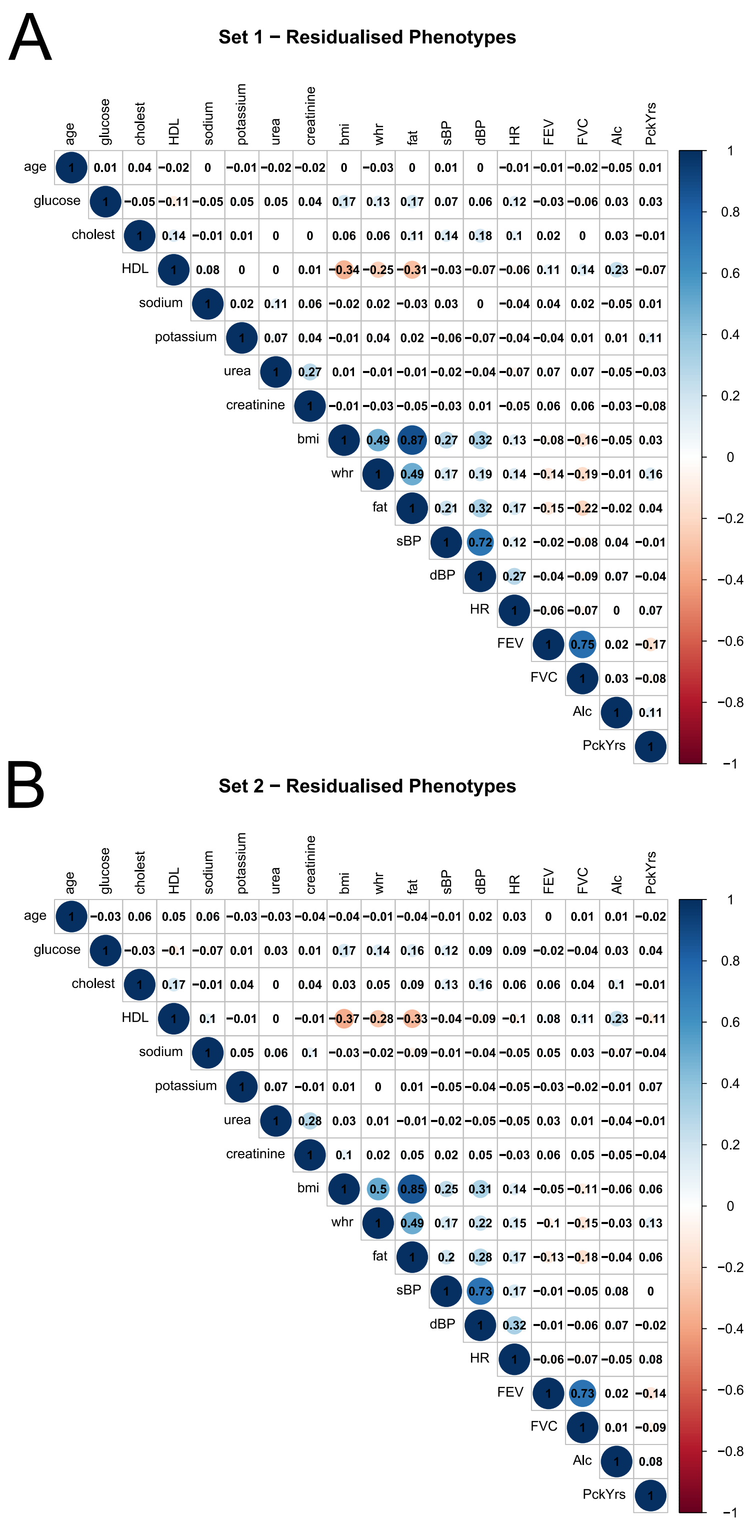


**Figure S2. Correlation structure between residualised phenotypes in Set 1 and Set 2 of Generation Scotland.** Set 1 (A) and Set 2 (B) had 2,578 and 4,450 unrelated individuals, respectively. Phenotypes were adjusted for chronological age and sex (and height for FEV and FVC). Age was not adjusted but is included for completeness of comparisons. Alc, alcohol consumption; bmi, body mass index; cholest, total cholesterol; dBP, diastolic blood pressure; fat, body fat percentage; FEV, forced expiratory volume in one second; FVC, forced vital capacity; HDL, high-density lipoprotein cholesterol; HR, heart rate; PckYrs, smoking pack years; sBP, systolic blood pressure; whr, waist-to-hip ratio

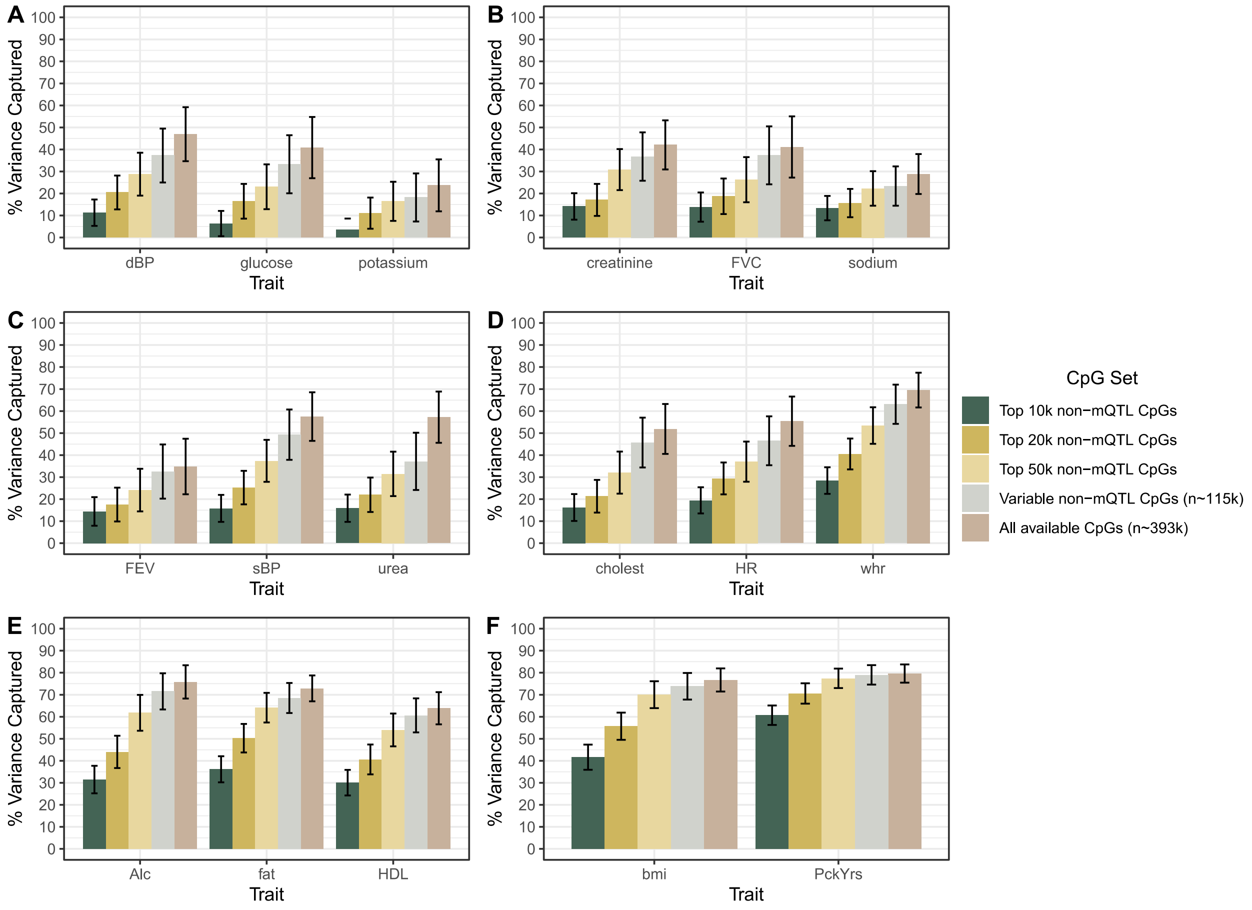


**Figure S3. Phenotypic variance captured by five nested sets of CpGs with decreasing numbers of CpGs and increasing mean variabilities.** Restricted maximum likelihood analyses were performed using blood DNAm and phenotypic data from 4,450 volunteers in Set 2 of Generation Scotland. Seventeen biochemical and complex traits are shown. The traits, or phenotypes, are ordered according to the amount of inter-individual variation captured by DNAm (smallest to largest). The seventeen traits are arranged into six groups (A – F). Vertical bars indicate 95% confidence intervals. Alc, alcohol consumption; bmi, body mass index; cholest, total cholesterol; dBP, diastolic blood pressure; DNAm, DNA methylation; fat, body fat percentage; FEV, forced expiratory volume in one second; FVC, forced vital capacity; HDL, high-density lipoprotein cholesterol; HR, heart rate; mQTL, methylation quantitative trait locus; PckYrs, smoking pack years; sBP, systolic blood pressure; whr, waist-to-hip ratio.

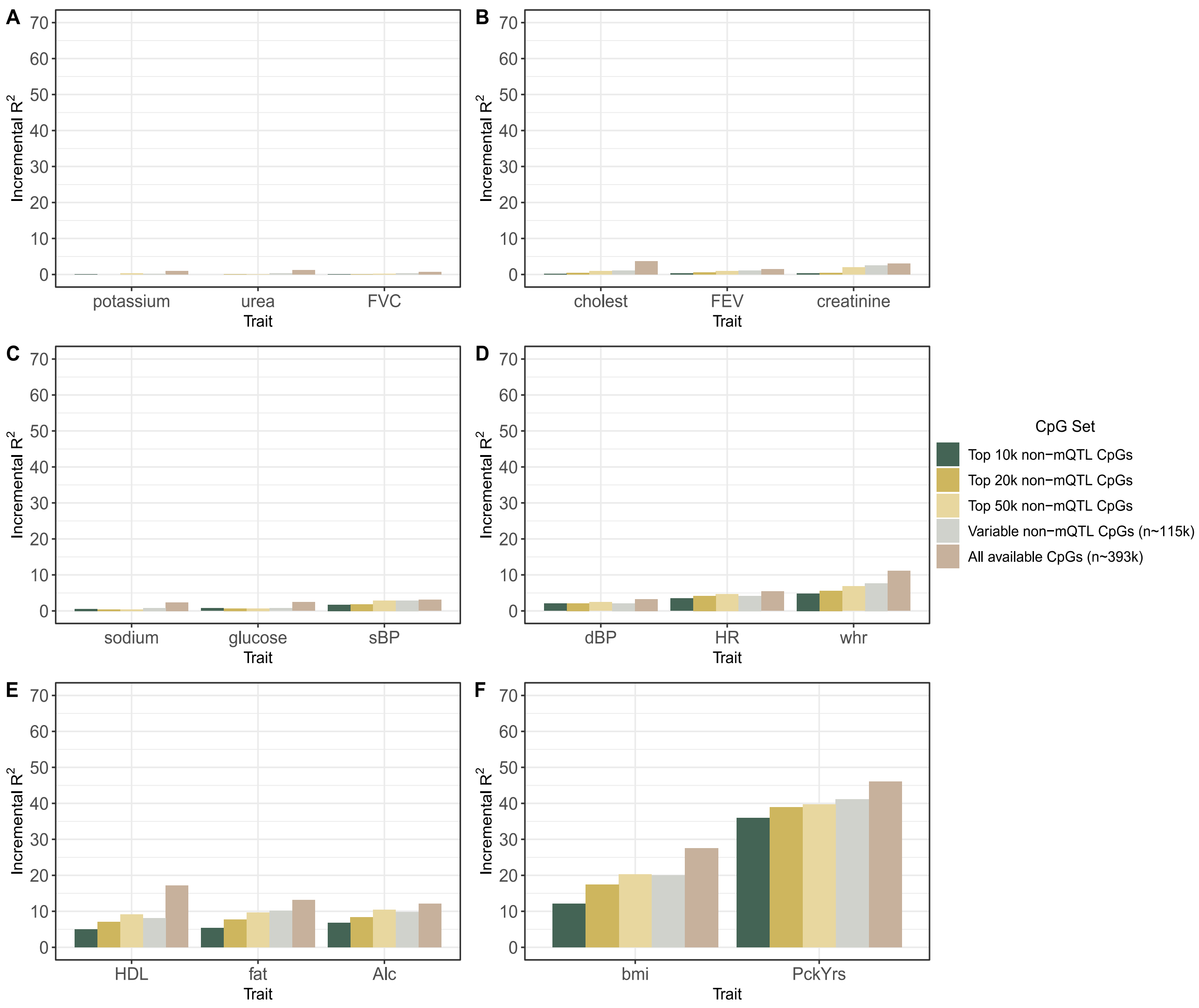


**Figure S4. Incremental $R^2$ estimates for DNAm-based predictors of seventeen traits using five nested sets of CpGs with decreasing numbers of CpGs and increasing mean variabilities.** LASSO regression was used to build DNAm-based predictors of seventeen traits using data from 4,450 volunteers in Set 2 of Generation Scotland. An unrelated sample of 2,578 individuals in Generation Scotland served as the test set. The traits, or phenotypes, are ordered according to the magnitude of incremental $R^2$ estimates (smallest to largest). The seventeen traits are arranged into six groups of three traits (A – F). Alc, alcohol consumption; bmi, body mass index; cholest, total cholesterol; dBP, diastolic blood pressure; DNAm, DNA methylation; fat, body fat percentage; FEV, forced expiratory volume in one second; FVC, forced vital capacity; HDL, high-density lipoprotein cholesterol; HR, heart rate; LASSO, Least Absolute Shrinkage and Selection Operator; mQTL, methylation quantitative trait locus; PckYrs, smoking pack years; sBP, systolic blood pressure; whr, waist-to-hip ratio.
